# Microbial N_2_O reduction in sulfidic waters: Implications for Proterozoic oceans

**DOI:** 10.1101/2025.06.28.662143

**Authors:** Steffen Buessecker, Aya S. Klos, Melanie E. Quan, Anna Finch, Morgan S. Sobol, Guy N. Evans, Ariel D. Anbar, Betül Kaçar, Anne E. Dekas

## Abstract

Throughout Earth’s history, shifts in ocean redox influenced the bioavailability of trace metals, shaping the activity of microorganisms. In Proterozoic oceans, the precipitation of copper (Cu) with sulfide was hypothesized to limit the bioavailability of Cu. This limitation may have suppressed microbial reduction of nitrous oxide (N_2_O), due to the Cu dependency of nitrous oxide reductase (Nos). It is thought that without this critical microbial sink, Proterozoic oceans were a significant net source of N_2_O. Here, we revisit this paradigm in light of recently derived ∼20-fold lower estimates for sulfide in Proterozoic seawater and an empirical evaluation of the potential for microbial N_2_O reduction under sulfidic conditions. Leveraging publicly available environmental metatranscriptomes, we infer active N_2_O reduction from the detection of *nosZ* transcripts in multiple marine and lacustrine systems in which sulfide and Cu concentrations are analogous to those of the Proterozoic. In controlled culture experiments, we demonstrate that the purple non-sulfur bacterium *Rhodopseudomonas palustris* can reduce N_2_O at sulfide concentrations up to 100 µM, well above levels predicted for Proterozoic oceans. Based on trace metal speciation modeling, we suggest that Cu remains bioavailable under Proterozoic-like conditions as a dissolved CuHS^0^ complex. Using phylogenetics, we infer that early N_2_O reducers were probably anoxygenic phototrophs and performed N_2_O reduction as dark metabolism. Collectively, these observations suggest microbial N_2_O reduction occurs under euxinic conditions, implying that Proterozoic marine N_2_O emissions were substantially lower than previously proposed. Our conclusions inform our understanding of the microbial ecology in sulfidic waters, the early climate, and the search for extraterrestrial life.

## Introduction

Trace metals in the environment impact global climate by catalyzing microbial metabolisms that control the biogeochemical cycles of greenhouse gases (Morel and Price, 2003). Integrated into the molecular tapestry of proteins, trace metals drive electron transfers important for the cycling of gases such as methane (CH_4_) and nitrous oxide (N_2_O). These impacts likely changed over geologic time because the redox evolution of the oceans profoundly affected the availability of trace metals to microbial cells (Anbar and Knoll, 2002; Anbar, 2008). For instance, the waning availability of nickel (Ni) in seawater may have affected biological formation of CH_4_ during the Archean eon, due to the dependence on Ni of methyl-coenzyme M reductase (Konhauser *et al*., 2009, 2015). Similarly, the presence of copper (Cu) in seawater influences the ocean’s capacity to act as a sink for N_2_O, due to the essential Cu cofactors of N_2_O reductase (Pomowski *et al*., 2011). Since atmospheric N_2_O has a radiative potential 300-fold that of carbon dioxide (Ravishankara *et al*., 2009), N_2_O accumulation arising from marine Cu scarcity could have induced a warmer Earth climate. This may have been particularly impactful when the Sun was fainter (Buick, 2007).

During the Proterozoic Eon, oceans likely contained orders of magnitude less Cu than today because the dissolved sulfide content of the oceans was higher under a low-O_2_ atmosphere (Saito *et al*., 2003). Sulfide has a high affinity to form precipitates with Cu, effectively stripping the trace metal from the water column. Prior considerations of Cu limitation of N_2_O reduction, which suggested Proterozoic seawater was a net source of N_2_O (Buick, 2007), were markedly influenced by the hypothesis that Proterozoic oceans were widely sulfidic (Canfield, 1998). However, research now suggests a more heterogenous ocean, mildly oxygenated at the surface, generally anoxic, and with extensive sulfidic wedges at the continental margins where high biological productivity overwhelmed oxidant supplies (Lyons *et al*., 2014; Fakhraee *et al*., 2025). Recent modeling of the sulfide concentrations in Proterozoic seawater indicates an upper limit of 2-3 µM (Ozaki *et al*., 2019), compared to >50 µM previously (Saito *et al*., 2003). Over time, as atmospheric O_2_ rose, hydrothermal Cu fluxes were increasingly complemented by riverine Cu input to the oceans stimulated by oxidative weathering. Subsequently, various inorganic and organic Cu ligands appear to have stabilized Cu in the water column (Stüeken, 2020; Zhao *et al*., 2016). The spatial variability in sulfide along with Cu complexes in potentially bioavailable forms open the possibility that Cu – and hence Cu-dependent microbial N_2_O reduction – was more prevalent than originally thought, rendering the ocean a sink for N_2_O rather than a source.

In this study, we combined environmental metatranscriptomics with microbial culturing and geochemical modeling to constrain microbial N_2_O reduction under sulfidic conditions. Globally mined metatranscriptomes included sequences from Proterozoic ocean analog sites of the Black Sea (2-30 µM sulfide) and Lake Cadagno (0.5-81 µM sulfide, Fig. 1). We assessed the limits of microbial N_2_O reduction in microbial cultures across a range of sulfide concentrations, and evaluated possible Cu source pools with a trace metal speciation model. To this end, we grew *Rhodopseudomonas palustris* as model organism for studying ancient metabolic processes (Larimer *et al*., 2004). We selected this purple non-sulfur bacterium as it was previously used to deduce trace metal interactions in the early ocean (Zhou *et al*., 2024). It also is a facultative phototroph, allowing us to investigate the photic zone as a potential habitat for early N_2_O-reducing microorganisms.

**Fig. 1.**
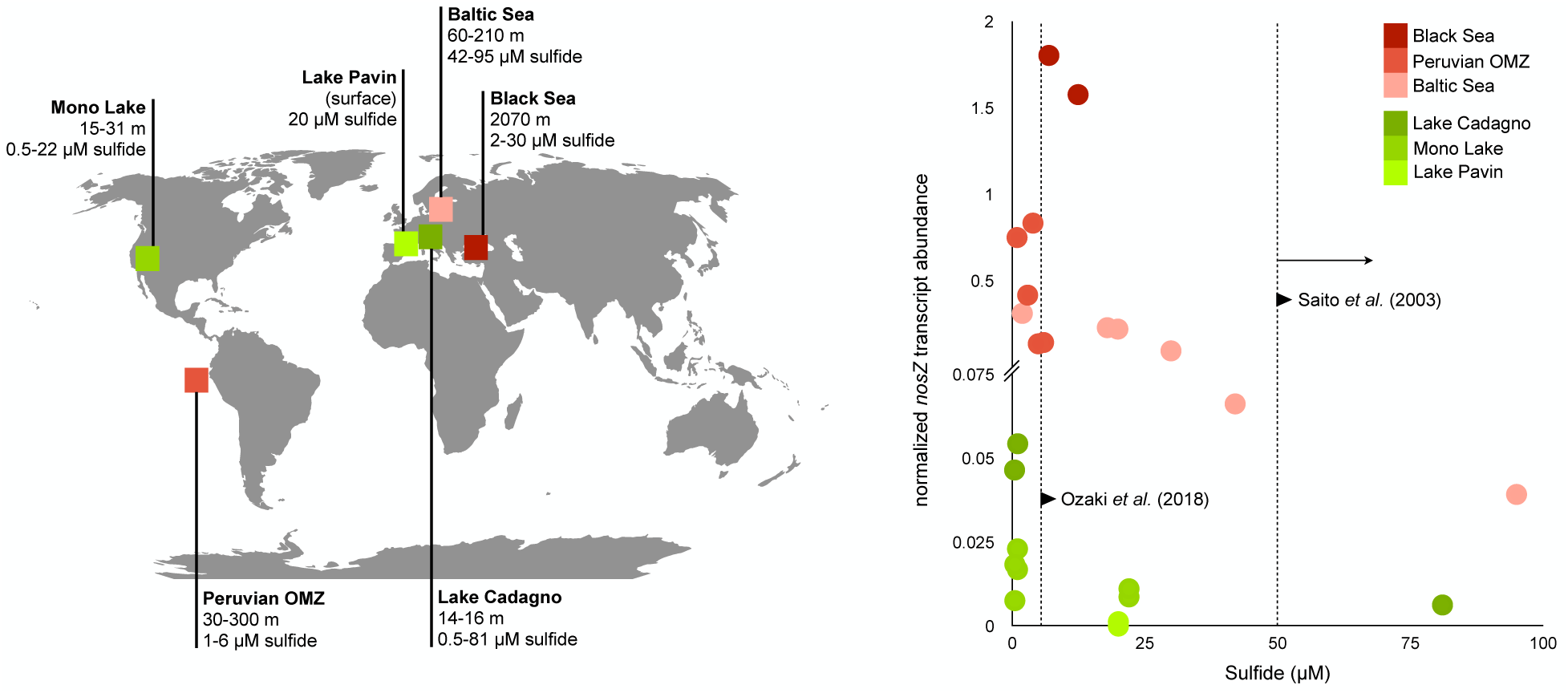
Normalized *nosZ* transcript abundance in metatranscriptomes of marine (red) and lacustrine (green) euxinia. Shown is the ratio of *nosZ* to *rplF* transcripts for comparison across datasets with different sequencing depths. Estimates of sulfide concentrations in Proterozoic oceans are shown with dotted vertical lines, with recent estimates in pelagic regions around 2-5 µM (Ozaki et al., 2018).

## Experimental Procedures

### Microbial culturing

*Rhodopseudomonas palustris* strain CGA009 (Niel, 1944) was anaerobically grown in modified liquid mineral medium (PM medium) with 20 mM sodium acetate as the carbon source and 0.01% ammonium sulfate as the nitrogen source. Oxygen was rigorously excluded from cultures and sulfide stocks through N_2_ sparging and storage in an anaerobic chamber. The medium was buffered with 500 mM Na_2_HPO_4_-KH_2_PO_4_ according to Oda and colleagues (Kim and Harwood, 1991; Oda *et al*., 2005). Cu was excluded from the trace metal solution (*metal 44*), though other media components supplied ∼30 nM of Cu, constituting a background concentration. Prior to incubations, all glassware was soaked consecutively with 2M HCl, 10 mM EDTA solution and deionized MQ (18.2 MΩ × cm) water. Borosilicate Balch tubes (27 mL) filled with 5 mL media under an ultra-high purity (>99.999%) N_2_ headspace contained with thick butyl rubber stoppers were inoculated with a 50 µL seed culture of *R. palustris*. Triplicate microcosms were placed into a static 30°C incubator at 15 cm distance to an incandescent light bulb to achieve 60 µmol m^−2^ s^−1^ light exposure. For dark conditions, the light was turned off. Sulfide amendments were prepared using a 5% sodium sulfide stock solution serially diluted in N_2_-sparged MQ water under strict exclusion of air. Negative (-) sulfide controls received anoxic MQ water only. Controls without inoculum at each sulfide level indicated no precipitation (that may alter the effective sulfide concentration) after they were allowed to sit overnight in the incubator prior to inoculation. Cell growth was approximated with optical density measurements at 680 nm using a Spectronic 20D+ (Thermo Scientific).

### Gas chromatography

Gas samples were collected by replacing headspace gas with N_2_ which was accounted for by a dilution factor. N_2_O concentration was measured with a Shimadzu GC-2014 equipped with a ^63^Ni electron-capture detector (ECD). Electron plasma was maintained at 1.5 nA at 325 °C. Separation was induced by a 1-m HayeSep N pre-column serially connected to a 5-m HayeSep-D column conditioned at 75 °C. Nitrogen gas (UHP grade 99.999%, Praxair Inc.) served as carrier at 21 mL min^−1^ flow rate. Calibration was conducted with customized standard mixtures (Scott Specialty Gases, accuracy ±5%).

### Inductively coupled plasma mass spectrometry (ICP-MS)

At the end of incubations, culture media from each Balch tube was filtered through nylon syringe filters (0.2 μm) and acidified with trace metal grade nitric acid (Fisher Chemical) to a final concentration of 2%. Aqueous Cu concentration was measured by inductively coupled plasma mass spectrometry (ICP-MS) on a Thermo Scientific iCAP RQ ICP-MS instrument. The calibrated analysis was verified by internal standards using reference solutions and blanks, following an established protocol (Stuckey *et al*., 2016).

### Trace metal speciation modeling

Thermodynamic modeling of aqueous Cu speciation, and that of Na, K, Zn, Fe, Mn, Co, Mg, Ca, and Ni present in the growth medium was performed using the React module of *The Geochemist’s Workbench* (Bethke, 2022). We used a modified database built on the packaged thermo.com.R7.tdat database, which is an expanded variant of the LLNL database that includes thermodynamic data for numerous transition metal organic complexes. Specific modifications include updated thermodynamic data for Cu-bisulfide complexes (Etschmann *et al*., 2010), and modified data for Zn-bisulfide complexes (Mei *et al*., 2016). Importantly, the relative species abundances we derive reflect a metastable state (what the microbial cells in our experiment are exposed to), rather than a long-term equilibrium.

The starting composition of the model is based on that of the growth medium, including the sodium acetate carbon source, phosphate pH buffer, and *metal 44* stock solution. Microbial cells may acquire Cu by excretion of organic chelators. Because of the general knowledge gap on these compounds, including stability constants, we did not account for them in our model. To mimic the addition of Na_2_S in the experiments, H_2_S and NaOH are iteratively added to the model in a 1:2 molar ratio to attain concentrations of 1000 µmol kg^−1^ and 2000 µmol kg^−1^, respectively. For consistency with ICP-MS data indicating that Cu precipitation was limited despite increased sulfide concentrations, and the lack of observed solid phases during the experiment, mineral precipitation was suppressed in the model configuration. The sulfide-sulfate redox pair was decoupled, reflecting the slow rate of sulfide-sulfate equilibration at ambient temperature (Ohmoto and Lasaga, 1982). All model components and concentrations used are provided in S1.

### Metatranscriptome analysis

Sequence data from microbial cDNA were mined from publicly available metatranscriptomes in the NCBI database selected from marine and lacustrine suboxic to anoxic water columns. We primarily included datasets that reported dissolved sulfide measurements at the time biomass was sampled for RNA extraction. The following metatranscriptomes were included in our analysis: ERR6496069, ERR6496070 (Black Sea); SRR1824297, SRR1824298, SRR1824300, SRR1824301, SRR1824302 (Peruvian OMZ); SRR11600252, SRR11600251, SRR11600250, SRR11600256, SRR11600257, SRR11600259 (Baltic Sea); SRR13495288, SRR13495289, SRR13495290 (Lake Cadagno); SRR3097676, SRR3097675, SRR3097678, SRR3097677, SRR3097682, SRR3097681 (Mono Lake); SRR17400026, SRR17400027, SRR17400028, SRR17400029 (Lake Pavin). A wider array of metatranscriptomes is listed in S1. Raw reads were assessed with FastQC (Brown *et al*., 2017) and trimmed with TrimAl v1.4.rev15 (Capella-Gutiérrez *et al*., 2009), requiring a min length of 30 bp and a minimum average read quality score of 25. Further trimming parameters were adjusted to ktrim = r, k = 23, mink = 11, hdist = 1, tbo qtrim = r. Trimmed reads were then reassessed with FastQC. High-quality reads were directly assembled to protein sequences with Plass (Steinegger *et al*., 2019) in default configuration, resulting in a minimum of 1.8 million sequences per sample. The protein sequences were then searched for NosL, NosD, and NosZ with GraftM (Boyd *et al*., 2018) and Kofamscan (Aramaki *et al*., 2020) based on trained profile hidden Markov models (HMMs; Eddy, 1998) for each gene. NosZ hits were manually verified by placing sequences on a tree to discern from cytochrome c oxidase subunit 2 (coxII).

### Phylogenetic reconstruction

We retrieved 7,458 prokaryote genomes from Web of Life (Zhu *et al*., 2019) after quality-filtering for completeness (>90%) and contamination (<5%) using CheckM2 (Chklovski *et al*., 2023). Reference NosZ protein sequences from *Stutzerimonas stutzeri* (clade I), *Anaeromyxobacter* sp. Fw109-5 (clade II), and *Wolinella succinogenes* (clade II), downloaded from UniProtKB on October 16, 2024, were used to search the genomes with BLASTp version 2.14.0 (Camacho *et al*., 2009), relying on an E-value threshold of 10^−5^, following methods used in Garcia et al. (2022). The reference sequences were chosen to represent the two known *nosZ* clades and an alternate form of clade II *nosZ* predominantly found in Epsilonbacteria that possesses a c-terminal monoheme cytochrome c domain (Simon *et al*., 2004). Next, BLASTp hits were annotated with KEGG Orthology (KO) identifiers based on a comparison with profile HMMs, using the KofamKOALA webserver with an E-value threshold of 10^-5^ (Aramaki *et al*., 2020). Top hits corresponding to K00376 (*nosZ*) were retained for further phylogenetic analysis which yielded 610 sequences in total. These sequences were further screened for 7 conserved histidine residues (Simon *et al*., 2004) and 5 conserved residues of the Cu_A_ domain (Savelieff *et al*., 2008) using a custom R script. Sequences missing at least one of these conserved residues or annotated as “cytochrome c” were removed from the dataset. Following previous methods (Garcia *et al*., 2022), the retained 595 NosZ protein sequences were aligned with MAFFT v7.505 (Katoh and Standley, 2013) using the -auto flag. The alignment was subsequently trimmed in TrimAl v1.4.rev15 (Capella-Gutiérrez *et al*., 2009) using the -automated1 option. The trimmed sequence alignment was then used to construct a maximum likelihood phylogenetic tree with IQ-TREE v2.1.4 (Trifinopoulos *et al*., 2016) based on 1000 ultrafast bootstraps and the best-fit model selected by ModelFinder (Kalyaanamoorthy *et al*., 2017). To include all complex mixture models of protein evolution in the model selection, we implemented the flags -mrate E,I,G,I+G,R and -madd C10,C20,C30,C40,C50,C60,EX2,EX3,EHO,UL2,UL3,EX_EHO,LG4M,LG4X,CF4. The final phylogeny was visualized using the ggtree package v3.12.0 (Yu *et al*., 2017) in R v4.4.1. We used the Oxyphen package (Jabłońska and Tawfik, 2019) to predict the oxygen status associated with select genomes. Briefly, this software assesses aerobic and anaerobic phenotypes based on the number of oxygen-processing enzymes and the presence of enzymes involved in secondary oxygen metabolism. Oxyphen predictions were additionally supplemented with information on oxygen tolerance collected from the Bacterial Diversity Database *BacDive* accessed on March 3, 2025 (Schober *et al*., 2024), as well as information obtained from the literature (S2).

Phototrophs among the taxa represented in our genome dataset were identified using metadata on trophic lifestyle (Hördt *et al*., 2020), supplemented by information obtained through a literature review (S2). The crystallized structure of clade II *nosZ* from *Pseudomonas stutzeri* (Pomowski *et al*., 2011) (PDB identifier: 3SBR) was downloaded from UniProtKB on March 17, 2025 and visualized in ChimeraX v1.8 (S3).

## Results

### NosZ gene expression in Proterozoic ocean analogs

We investigated and detected transcripts of *nosZ,* the functional marker gene of N_2_O reductase, in metatranscriptomes from 6 modern euxinic systems (3 marine and 3 lacustrine; Fig. 1). Total sulfide abundances were measured at the time of RNA sampling, allowing us to directly relate expression of the *nosZ* gene to sulfide abundance. The metatranscriptomic data show *nosZ* transcript levels at sulfide concentrations up to nearly 100 µM (data at higher sulfide concentrations was not available), with highest expression below 25 µM—still well above that most recently predicted for the pelagic Proterozoic ocean. We additionally observed two prevailing trends. First, *nosZ* transcript count decreased with increasing sulfide concentration. This drop was particularly apparent in marine systems where a 10-fold increase in sulfide was accompanied by a 10 to 40-fold decrease in *nosZ* transcripts (Fig. 1). Second, normalized transcript counts from marine systems were about an order of magnitude higher than those from lacustrine systems. Within these two groups, highest *nosZ* expression was captured in the sulfidic waters of Proterozoic ocean analog Lake Cadagno (Philippi *et al*., 2021) and from a depth of 2070 meters in the Black Sea water column (Henkel *et al*., 2022). Here, *nosZ* transcripts were almost twice as abundant as *rplF* transcripts, a housekeeping gene we chose to normalize the transcript frequencies.

We refined our investigation into the transcriptional activity of the N_2_O reductase complex (Nos) by examining the catalytic site of derived structures from select *nosZ* sequences, and by searching for *nosL*, *nosD*, and *nosZ* transcripts in a wider array of metatranscriptomes. Microbial cells actively scavenge Cu from the environment using the membrane-anchored chaperone NosL. Once Cu is retrieved, its insertion into the CuZ center is facilitated by a conformational interaction between NosL and NosD, ensuring the maturation of functional NosZ (Müller et al., 2022). The additional OMZ sites and hydrothermal vent plumes are known sulfidic systems but sulfide measurements were not as closely coupled to the metatranscriptomes. We therefore could not associate transcript levels to distinct sulfide concentrations as done above.

We inspected the protein structure derived from Alphafold2 (Jumper *et al*., 2021), decorated it with cofactors by Alphafill (Hekkelman *et al*., 2023), and confirmed the expected Cu atoms coordinated by sulfur moieties (S4). The genes nosL and nosD for the accessory proteins aiding in Cu retainment are co-located to *nosZ* in all examined genomes (S5). They also showed transcription patterns consistent with those of *nosZ* in 47 metatranscriptomes retrieved from hydrothermal vent plumes (S1). In contrast, in 36 metatranscriptomes from OMZs, *nosL* was less transcribed than *nosD* and *nosL* by at least one order of magnitude. Nevertheless, our refined sequence assessment demonstrated that the *nosZ* transcripts detected in euxinia are indeed dependent on Cu and that the accessory proteins driving Cu insertion into N_2_O reductase are also expressed, with the exception of *nosL* in OMZs.

### Active N_2_O reduction up to 100 µM sulfide concentration

We exposed *R. palustris* to sulfide concentrations of 0, 10, 100, and 1000 µM to place constraints on the threshold at which N_2_O reduction ceases. We define the reported sulfide as the sum of all sulfide species added. In dark incubations, N_2_O was completely consumed after 66 hours at sulfide concentrations of up to 100 µM (Fig. 2a). N_2_O was not consumed for the duration of the dark experiment at 1000 µM sulfide. There was no significant difference in the overall N_2_O reduction rate at 10 to 100 µM sulfide (0.14-0.2 ppm hour^−1^, *p* < 0.05). We also simulated permanent day conditions to reveal light response. While without sulfide dark cultures reduced 85% of N_2_O at the ∼40 hour mark, only 23% of N_2_O was reduced by light-exposed cultures. Notably, N_2_O reduction was retarded in the presence of 10-100 µM sulfide in light relative to the same sulfide range in dark incubations. *R. palustris* photosynthesized during light exposure based on production of pigments absorbing strongly at 680 nm (Fig. 2b). To assure the observed N_2_O effects were not due to reduced biomass growth, we used optical density at that wavelength as proxy for cell growth and determined no relevant negative biomass effects, even at the highest sulfide concentration (1000 µM).

**Fig. 2.**
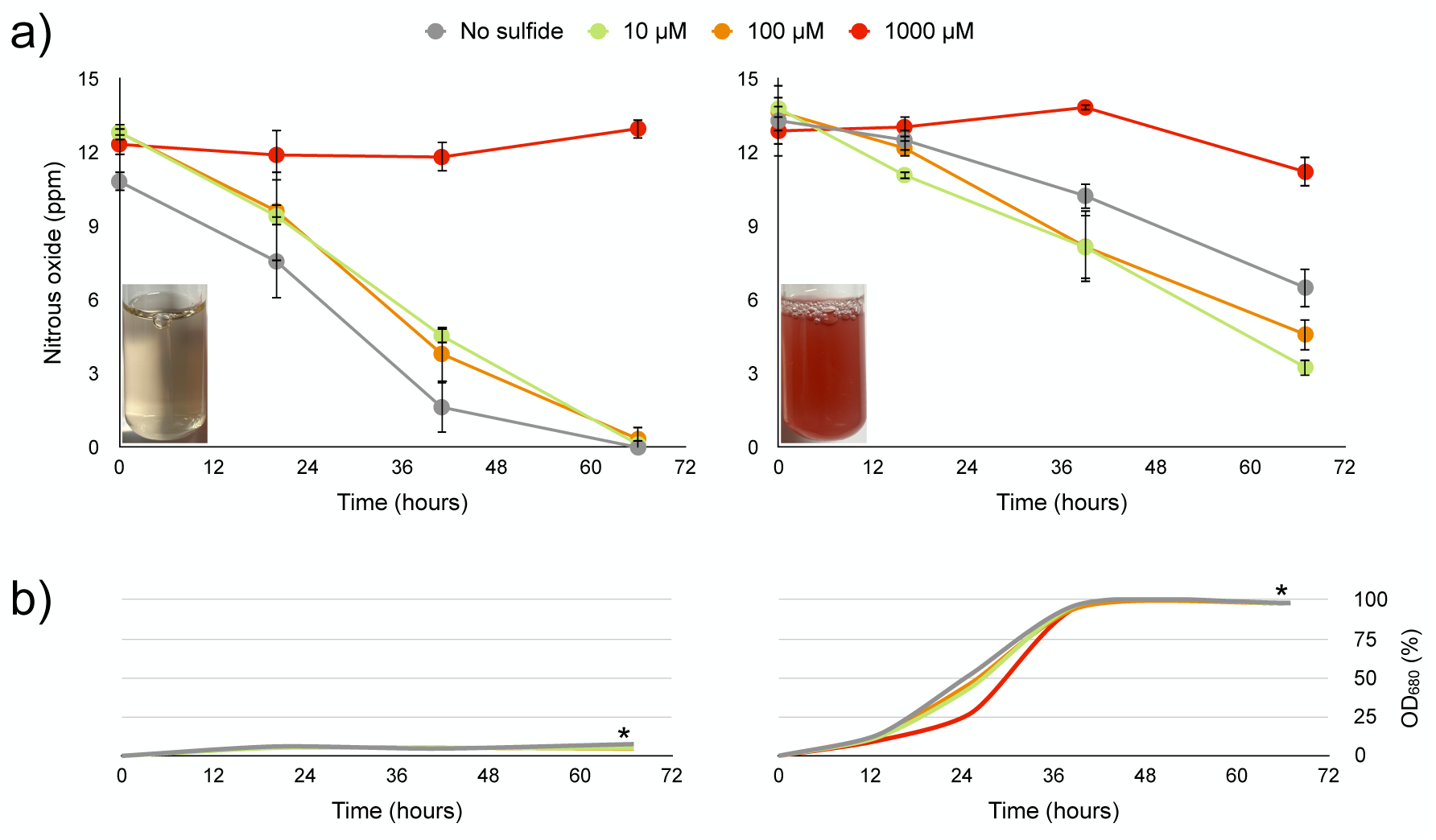
N_2_O reduction in *Rhodopseudomonas palustris* cultures. Headspace N_2_O (**a**) and cell growth measurements (**b**) during continuous dark (left) or light (right) conditions. The culture was incubated at different sulfide concentrations (0-1000 µM, n=3). Photographs show response in pigment production and were taken at the time point marked by asterisks.

### Cuprous hydrosulfide (CuHS) is the dominant inorganic sulfur species under physiological sulfidic conditions

To explore the bioavailability of Cu in the culture, we measured total dissolved Cu with ICP-MS and modeled Cu speciation along a sulfide gradient (dark conditions only). Our method accounts for total dissolved Cu in the <0.2 μm fraction after filtering and acidification, including all soluble Cu species, such as Cu^2+^, Cu^+^, and Cu–ligand complexes like CuHS^0^ and Cu(HS)_2_^−^, which are converted to free Cu ions during acid digestion. Media without inoculum contained ∼30 nM Cu. Cultures with 0 uM, 10 µM, 100 µM, and 1000 µM sulfide treatments contained 28.9±2.3 nM, 20.3±2.1 nM, 16.7±0.3 nM, and 13.1±0.5 nM of total dissolved Cu, respectively. We presume the 3-27% difference between inoculated and uninoculated media is accounted for by cellular uptake.

To evaluate the Cu species present at the time of active N_2_O reduction, we constrained the model to reflect metastable aqueous speciation rather than long-term equilibrium. Thus, because we did not observe precipitate formation within the time of the experiments, all thermodynamic calculations were performed by suppressing mineral precipitation of chalcocite (Cu_2_S) and pyrite (FeS_2_). These are both supersaturated in equilibrated sulfide-bearing solutions. We modeled Cu speciation based on the media composition (S6) at different sulfide levels. Freely dissolved Cu^+^ (the dominant redox state in reduced sulfidic milieu) is only present when sulfide levels are below 10 µM. In this range, the cupric phosphate complex Cu(PO_4_)^−^ is predicted to constitute a minor component of dissolved Cu species (Fig. 2). The dominant Cu species is the bisulfide complex CuHS^0^ below 250 µM total sulfide, and Cu(HS)_2_^−^ above that concentration. Although acetate is 10-10^3^ times more concentrated than sulfide, it constitutes a negligible organic ligand to Cu because the stability constant of the CuHS^0^ complex is ∼10^10^ times that of the CuAc complex (Mountain and Seward, 1999).

### Microbial evolution of N_2_O reduction

To extend our investigation of N_2_O cycling into deep-time, we conducted a phylogenetic analysis of N_2_O-reducing microbes. N_2_O reductase is the key enzyme enabling microbial N_2_O reduction and is phylogenetically classified into two distinct Cu-dependent clades (Hallin et al., 2018). Based on 595 highly conserved *nosZ* sequences identified in microbial genomes obtained from the Web of Life reference phylogeny, we evaluated taxonomy, oxygen status, phototrophy, and presence of nitrite reductases (Fig. 4). Phyla encoding Clade I *nosZ* comprised canonical proteobacterial denitrifiers as well as novel Euryarchaeota, whereas those with Clade II *nosZ* were more diverse, including Bacteroidota, Bacillota, Verrucomicrobiota, and Chlorobiota. Archaea with previously unrecognized N_2_O-reducing potential were primarily affiliated to halophilic groups (*Halobacteriales* and *Natrialbales*) typically encountered in low-oxygen brines. We categorized the N_2_O-reducing genomes into aerobic, facultative, microaerophilic, and anaerobic using marker enzymes typically involved in oxygen metabolism. The majority of aerobic genomes were found to encode Clade I *nosZ*. Anaerobic genomes were aggregated in a cluster of Pseudomonadota within Clade II, although a large number of that clade could not be characterized by oxygen status due to ambiguous Oxyphen classifications and unavailable data for certain taxa in *BacDive*. Most of these anaerobe-associated genomes with clade II *nosZ* belonged to Epsilonproteobacteria, even though Alpha-, Beta-, Gamma-, and Deltaproteobacteria were included as well. Of 287 bacterial genera (32 genomes without genus affiliation), 19 were potential phototrophs. Of these 19, three (*Rhodopseudomonas, Roseobacter, and Rubrivivax*) were facultative anaerobes and two (Chlorobi bacterium and *Thiocapsa*) were anaerobes, pointing to modern phototrophic N_2_O reducers as mainly aerobes.

Cu-dependent nitrite reductases catalyze the one-electron reduction of nitrite to the gaseous species nitric oxide (NO). The nirK gene encodes a cupredoxin-like domain to assemble a Cu-containing active site, while the nirS gene contains an iron center instead. Using one form of nitrite reductase over the other may point towards the environmental availability of Cu. Expectedly, 276 of all 595 N_2_O -reducing genomes also possessed *nirK*, while only 127 possessed *nirS*. Clade II genomes in particular showed *nirK* genes and no *nirS* genes, underlining their dependence on Cu metalloenzymes (Fig. 4).

## Discussion

Our findings suggest that microbial N_2_O reduction was feasible under conditions estimated for Proterozoic oceans. We observed substantial *nosZ* transcription across multiple euxinic marine and lacustrine systems, suggesting microbes can reduce N_2_O within natural euxinic waters exceeding the sulfide concentrations estimated for the Proterozoic (Fig. 1). Both the Black Sea and Lake Cadagno are commonly considered analogous to Proterozoic conditions due to the persistence of anoxia and stability of the redoxclines, and exhibited highest *nosZ* transcript numbers among marine and lacustrine systems, respectively. Of note, the Black Sea contains total dissolved Cu concentrations of 0.2-2 nM (Luther *et al*., 1991), similar to estimates for the Proterozoic (Stüeken, 2020). Unfortunately, Cu concentration data for Lake Cadagno or the other euxinic systems were scarce. The assumption that detectable *nosZ* transcripts are indicative of an active N_2_O reduction pathway is not definitive. However, previous culture experiments have indeed shown a positive correlation between *nosZ* expression and N_2_O depletion, indicating a functional coupling between gene expression and enzymatic activity (Sullivan *et al*., 2013; Liu *et al*., 2014). The generally higher transcript levels in marine systems may be explained by higher production of organic substrate for N_2_O reducers compared to more oligotrophic lakes (Ward *et al*., 2008).

Our culture experiments indicate that N_2_O reduction activity at high (∼100 µM) sulfide concentrations (Fig. 2) is likely explained by microbial sourcing of Cu from dissolved Cu– sulfide complexes, such as CuHS^0^ (Fig. 3). While in previous microcosm experiments microbial N_2_O reduction was impaired when marine denitrifying cultures were starved of Cu (Granger and Ward, 2003; Manconi *et al*., 2006; Felgate *et al*., 2012), effects of distinct sulfide levels on microbial N_2_O reduction have not been explored. Moreover, the Proterozoic seawater dominant species CuHS^0^ (Stüeken, 2020) and organic Cu complexes, such as coordinated with thiols glutathione and cysteine (Leal and Berg, 1998), were potentially bioavailable for early organisms (Zhao *et al*., 2016). Our model organism *R. palustris* consumed N_2_O in culture incubations amended with 100 µM sulfide, which exceeds probable sulfide levels of Proterozoic seawater (Saito *et al*., 2003; Ozaki *et al*., 2019). At those sulfide levels, 80-100% of Cu was present as CuHS^0^ (Fig. 3). The demise of N_2_O reduction between 100 and 1000 µM sulfide (Fig. 2) is consistent with a shift in the Cu speciation from CuHS^0^ to Cu(HS)_2_^−^ (Fig. 3). Thus, we demonstrate for the first time that a Cu-dependent microbial metabolism can utilize Cu bound in a sulfidic complex, a form likely abundant on the early Earth.

**Fig. 3.**
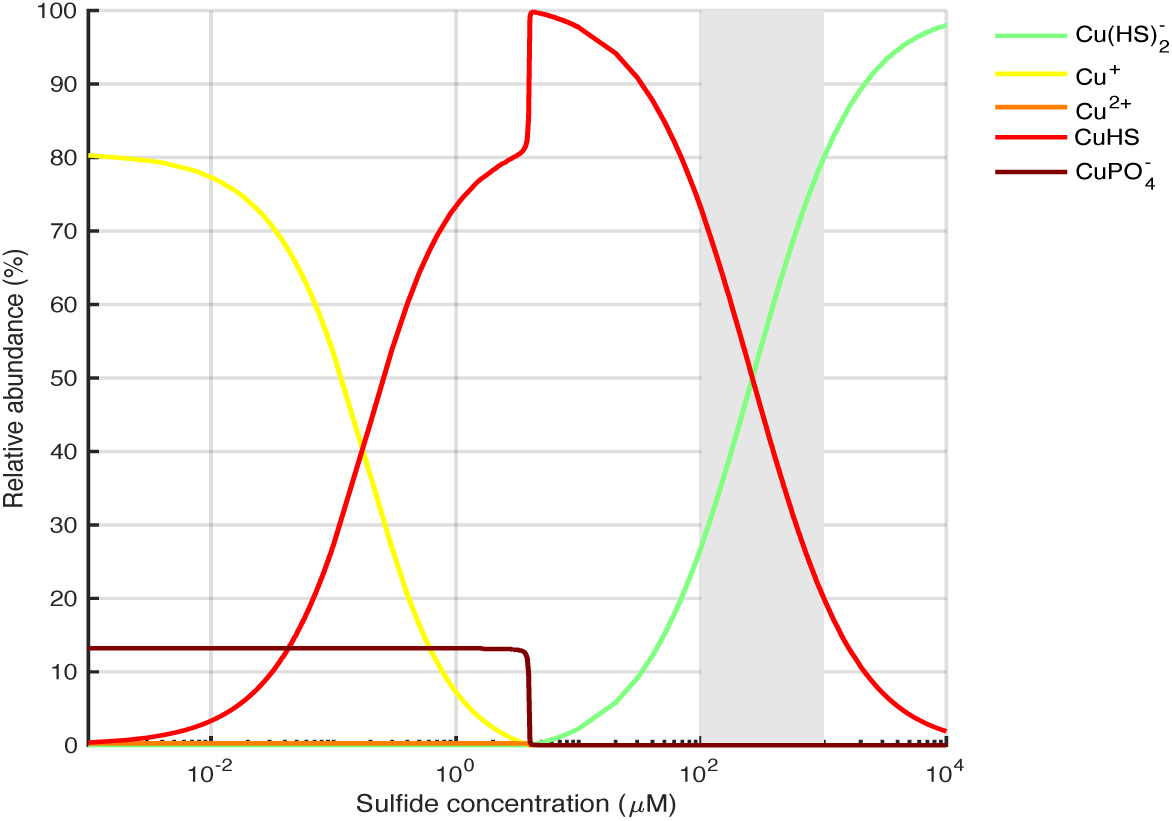
Cu speciation model output. Gray shading marks region within which an inhibition of microbial N_2_O reduction in *R. palustris* occurred. Organic acid complexation with known organic compounds was found to be negligible. While organic ligands generally increase the solubility of Cu, these would need to be specialized to effectively compete with sulfide.

The dominance of Cu as CuHS^0^ likely has biological consequences. As a neutral species, CuHS^0^ has the advantage of being able to passively penetrate the outer cell membrane, owing to the plasticity of the fatty acid bilayers that compose bacterial membranes in sulfidic waters (Boukhchtaber *et al*., 2025). We postulate that when CuHS^0^ dissociates into Cu^+^ and HS^−^, potentially due to the pH change during migration from the extracellular environment into the periplasm, the monovalent Cu ion can be readily chelated by NosL. It is conceivable that other chelators act on intracellular Cu^+^ or that sufficient Cu concentration allows for a direct uptake by the ATP-binding-cassette complex NosDFY (Müller *et al*., 2022). This would explain the observed anomaly in *nosL* expression in OMZ metatranscriptomes (S1). Irrespectively, the dominant CuHS^0^ species appears to be a viable Cu source for N_2_O reductase. In light of this evidence, further research on the molecular-scale Cu acquisition from CuHS^0^ is needed to characterize the underlying biochemical mechanism.

We propose that the marine photic zone could have constituted a sustained N_2_O sink fueled by phototrophic activity during the Proterozoic (Fig. 4). While modern marine N_2_O-reducing microbes are predominantly heterotrophs that oxidize organic substrate with N_2_O in OMZs (Intrator *et al*., 2024), it is possible that early N_2_O reducers were more often facultative photoautotrophs. The ability to fix CO_2_ would have made them less dependent on exogenous organic carbon. Diverse anaerobic phototrophs are capable of N_2_O reduction (McEwan *et al*., 1985). Anaerobic N_2_O reduction is coupled to ATP generation, thereby providing an alternative energy source for these phototrophs under dark conditions (McEwan, 1994). Thus, dark N_2_O reduction could yield a competitive advantage during day-night cycles within the marine photic zone – a potentially extensive N_2_O-consuming microbial habitat of the pelagic ocean in the Proterozoic.

**Fig. 4.**
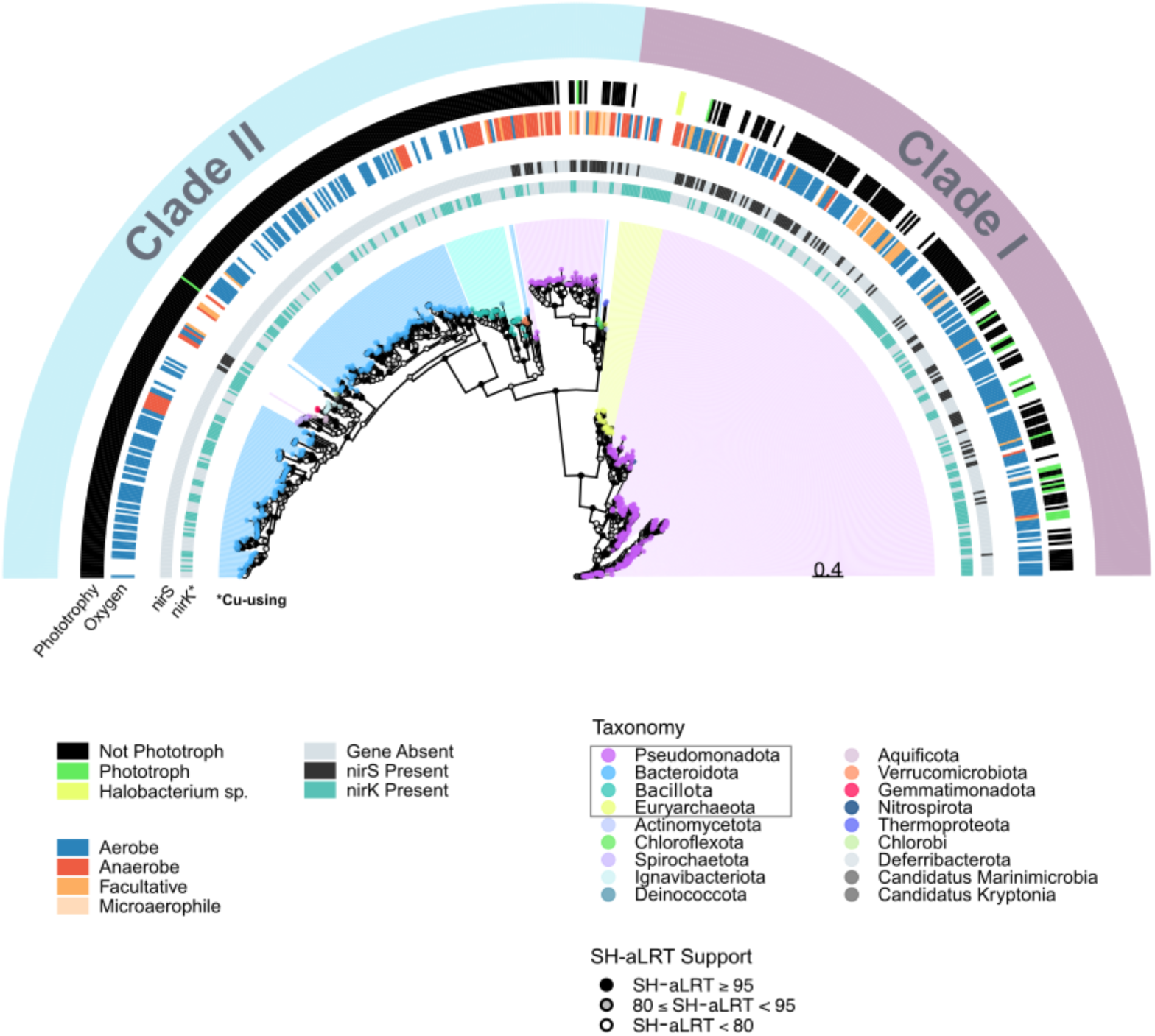
Unrooted maximum likelihood phylogenetic tree of NosZ protein sequences. The phylogeny includes 595 *nosZ* sequences collected from genomes obtained from Zhu *et al*. (2019). Clade I (purple) and clade II (blue) *nosZ* sequences show distinct clade separation in the tree. Branch tips are colored by taxonomy at the level of phylum. Four phyla (Pseudomonadota, Bacteroidota, Bacillota, and Euryarchaeota—boxed in the legend) forming large, notable clades within the tree are further distinguished by shading beyond branch tips. Metadata is shown for specific taxa with genomes represented in the tree. These include, phototrophy (phototrophs vs non-phototrophs), oxygen status (aerobes, anaerobes, facultative organisms, and microaerophiles), and presence or absence of nitrite reductase genes *nirS* and *nirK* in the corresponding genomes. Halobacterium species are distinguished within the “phototrophy” category due to previous characterization of photobiology within the aerobic archaeon *Halobacterium salinarum* NRC-1 (DasSarma *et al*., 2001). Note that *nirK*, like *nosZ*, is a Cu-using enzyme and is distinguished in the tree.

As observed in modern OMZs, dissolved Cu could have supported an anaerobic/microaerophilic nitrogen cycle (Glass *et al*., 2015) linked to organic carbon production by photosynthesis (Fig. 4). Nitrification could have served as a biotic N_2_O source (Casciotti *et al*., 2003; Santoro *et al*., 2011) and was probably complemented by abiotic N_2_O fluxes from aqueous Fe^2+^ (Stanton *et al*., 2018) and reduced Fe minerals (Buessecker *et al*., 2022) in Fe-rich parts of the ocean (Planavsky *et al*., 2011). Regardless of whether N_2_O was reduced directly by phototrophs or used by heterotrophs that benefited from photosynthetically fixed carbon, substrate dynamics likely obliged day-night cycles. Microbial cells may have fixed inorganic carbon via photosynthesis during the day and respired organic carbon using N_2_O at night. Accordingly, marine N_2_O emissions would have contributed less to warming the Proterozoic climate than estimated based on Cu-scarce conditions (Buick, 2007; Roberson *et al*., 2011; Stanton *et al*., 2018). Our findings support the concept of a shared evolutionary history between denitrification enzymes (Parsons *et al*., 2021) and protein complexes involved in aerobic respiration (David and Alm, 2011; Stanton *et al*., 2018). The photic zone, serving as both a habitat for N_2_O reducing microbes and a hotspot for emerging oxygenic photosynthesis, brought N_2_O and oxygen respiration into direct ecological proximity. This convergence likely promoted biochemical interactions that shaped the evolutionary trajectory of cellular respiration.

Taken together, these insights prompt a reassessment of the long-standing assumption that Proterozoic oceans were a major source of atmospheric N₂O, as microbial N₂O reduction appears to have been biogeochemically viable during this period. This reevaluation carries important astrobiological implications: On early Earth, significant accumulation of marine-derived N_2_O likely required elevated H_2_S concentrations (>100-1000 µM), suggesting that the detection of N_2_O in an exoplanetary atmosphere may depend on specific redox and biogeochemical conditions. These factors need to be considered in future life-detection strategies that rely on atmospheric N_2_O as a potential indicator of biological activity (Schwieterman *et al*., 2018, 2022).

## Acknowledgments

We thank the scientific community for field sampling, generation of metatranscriptome data, and public deposition of all data sets used in our study. We also thank Karen Casciotti for help with the interpretation of gene transcription results. We acknowledge the Environmental Measurements Facility (EMF) at the Stanford Doerr School of Sustainability for liquid and gas phase analyses, and the Center for High Throughput Computing (CHTC) at the University of Wisconsin–Madison for providing computing resources. This work was supported by the National Aeronautics and Space Administration (NASA) Interdisciplinary Consortium for Astrobiology Research: Metal Utilization and Selection Across Eons, MUSE (Grant no. 80NSSC17K0296). S.B. and M.S. were supported by the NASA Postdoctoral Program, administered by Oak Ridge Associated Universities under contract with NASA. A.S.K. was supported by the Paglia Post-Baccalaureate Research Fellowship from Carleton College.

## Data Availability Statement

The manuscript and supplementary material together provide all data necessary to evaluate the study’s conclusions. Raw data shown in the plots can be requested from the authors.

## Supplementary

**Supplement 1.**
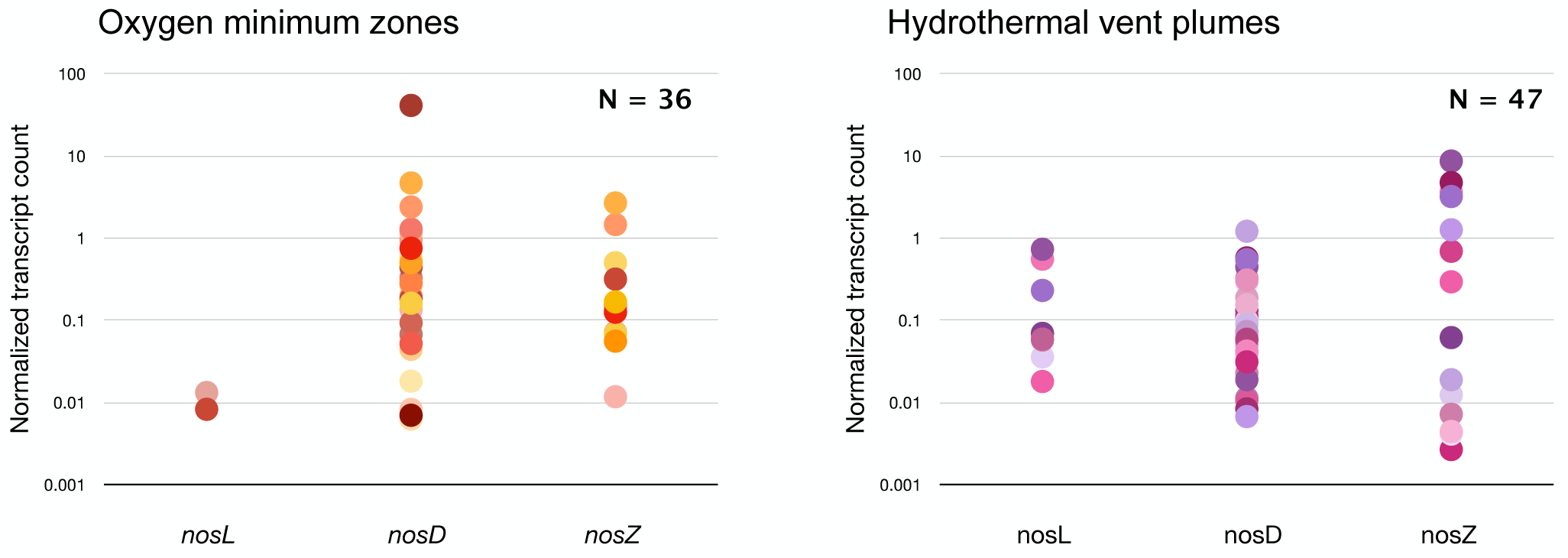
*nosL*, *nosD*, and *nosZ* transcriptional activity in microbial communities of OMZs and hydrothermal vent plumes. The wider array of metatranscriptomes included ERR6496070, SRR1824297, SRR1824298, SRR1824300, SRR1824302, SRR1839865, SRR1839866, SRR1839868, SRR1918203, SRR3001182, SRR3001183, SRR3001186, SRR3001187, SRR3001188, SRR3001190, SRR3028162, SRR3710727, SRR3710729, SRR3719474, SRR3732357, SRR3732359, SRR3732360, SRR3882725, SRR5204523, SRR5264433, SRR5264438, SRR5264512, SRR5264513, SRR5264545, SRR5264546, SRR5264547, SRR5264548, SRR5909415, SRR5909421, SRR6214539, SRR9696254 (representing OMZs), and SRR24080074, SRR24080075, SRR24080076, SRR24080077, SRR24080078, SRR24080079, SRR24080080, SRR24080091, SRR24080113, SRR24080135, SRR24080169, SRR24080171, SRR24080173, SRR24080175, SRR24080177, SRR24080178, SRR24080180, SRR24080182, SRR24080184, SRR24080242, SRR26701688, SRR26701692, SRR26701693, SRR26701694, SRR26701695, SRR26701696, SRR26701697, SRR26701699, SRR26701700, SRR26701704, SRR26701706, SRR26701709, SRR26701710, SRR26701712, SRR26701713, SRR26701714, SRR26701715, SRR26701716, SRR26701731, SRR26701732, SRR26701733, SRR26701734, SRR26701735, SRR26701736, SRR26701737, SRR26701738, SRR26701739 (representing HVs).

**Supplement 2.** ∼in accompanying Excel file∼.

**Supplement 3.**
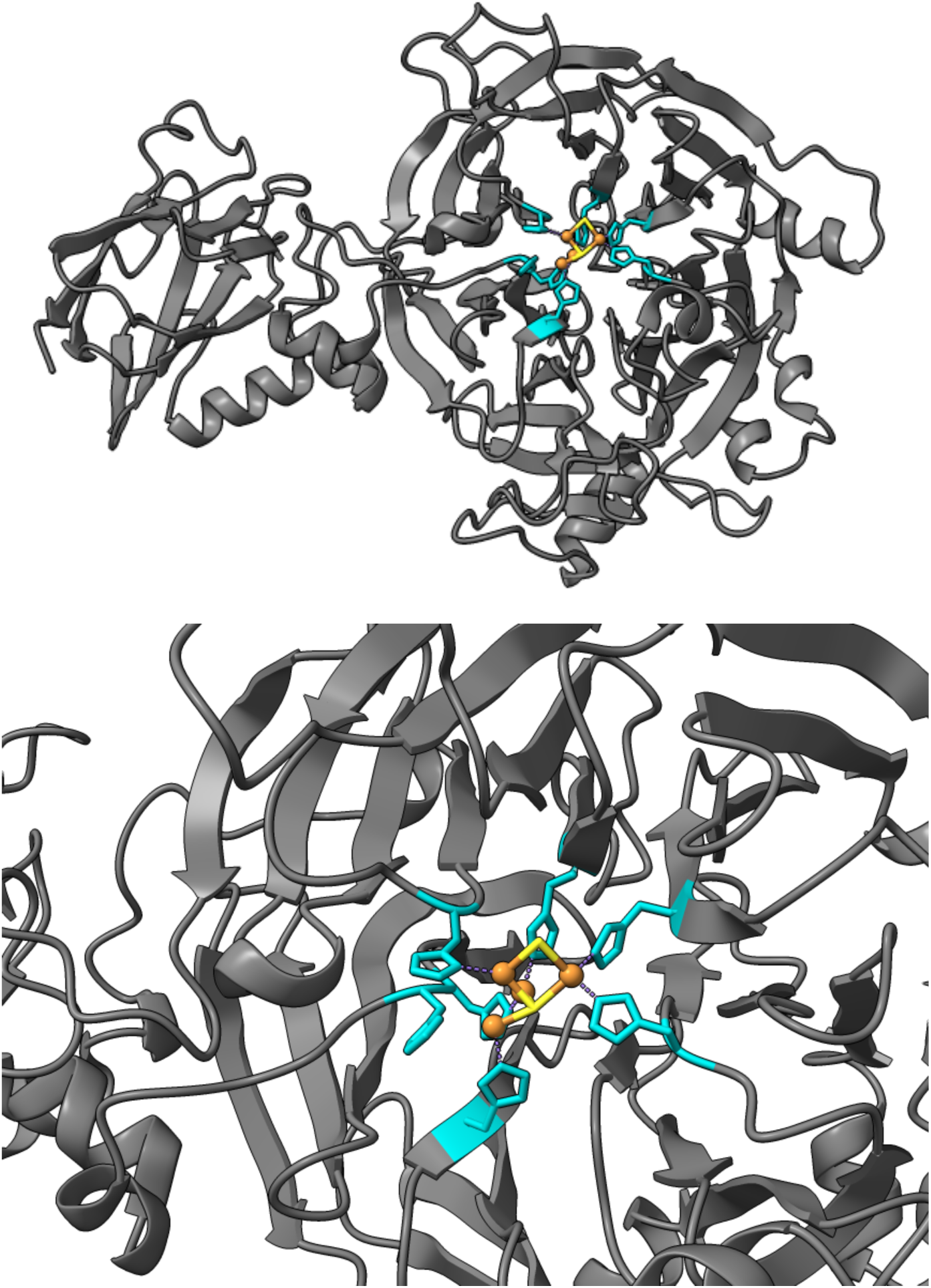
Visualization of *Pseudomonas stutzeri* nosZ (clade II) based on the crystal structure identified by Pomowski *et al*. (2011); PDB ID: 3SBR. Only one subunit of the NosZ heterodimer is shown (top). A zoomed in image of the CuZ center in the *nosZ* active site is shown (bottom). Histidine residues coordinating the *nosZ* CuZ center are highlighted in turquoise where interactions are shown as purple dotted lines. Cu atoms are shown in orange while bonds with coordinating sulfur atoms are shown in yellow.

**Supplement 4.**
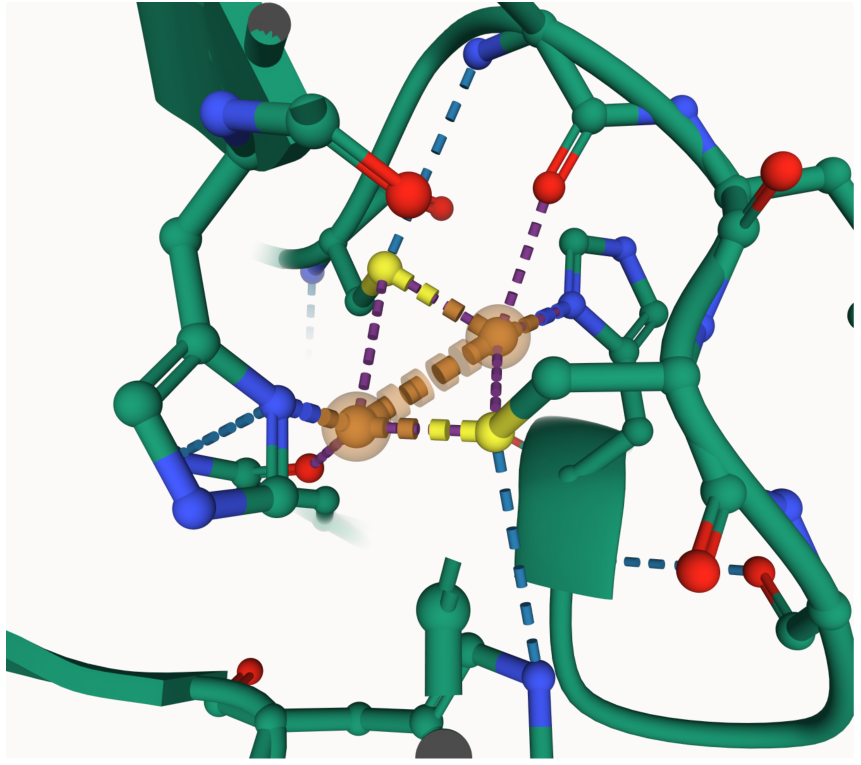
Alphafill-decorated model of the catalytic CuZ site in N_2_O reductase indicating 2 Cu atoms (orange) to match the topological cavity.

**Supplement 5.**
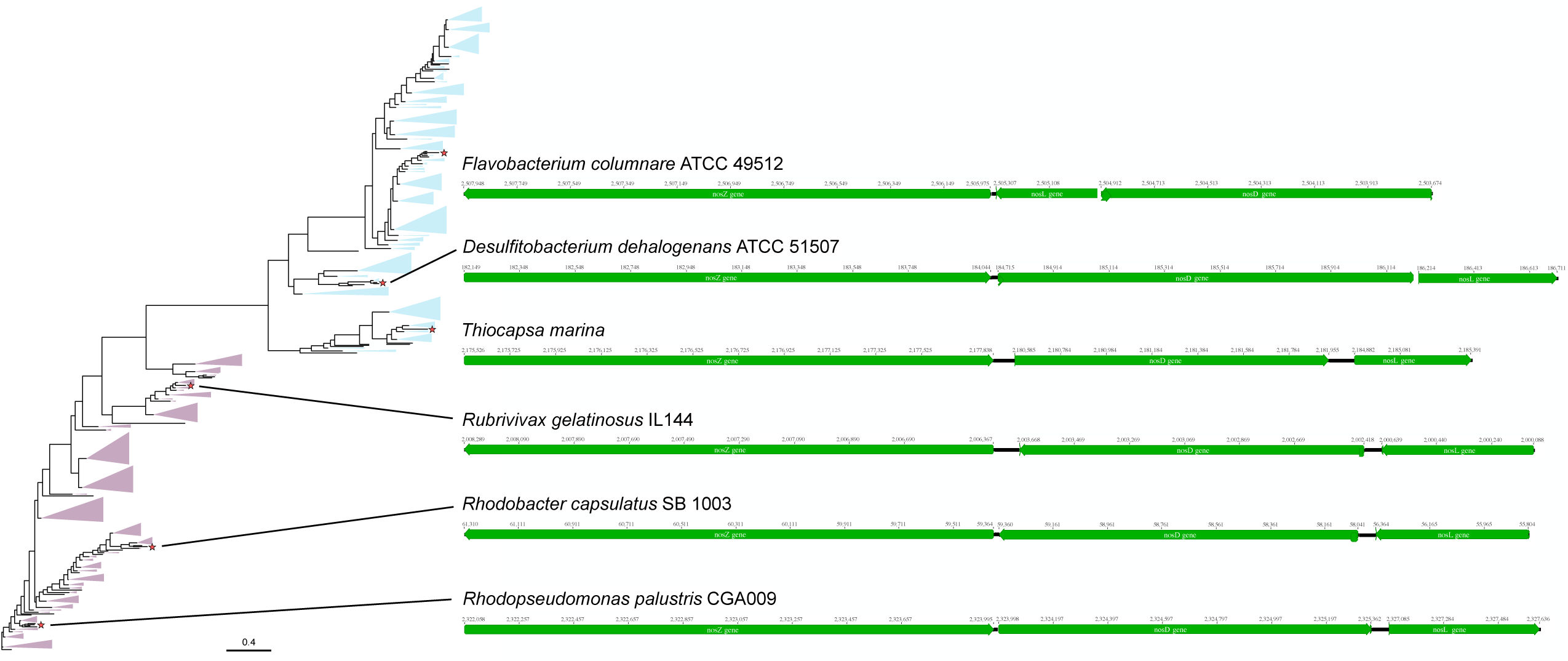
Comparison of *nos* gene clusters in six select clade I and II genomes.

**Supplement 6.** Model components and respective abundances.

| **Component** | **Concentration** |
| --- | --- |
| pH | 6.8 |
| Na<sup>+</sup> | 45 mM |
| HPO<sub>4</sub><sup>--</sup> | 25 mM |
| K<sup>+</sup> | 12.5 mM |
| Lactic Acid | 2 mM |
| Mg<sup>++</sup> | 29 μM |
| SO<sub>4</sub><sup>--</sup> | 7 μM |
| Zn(II) | 3809 μM |
| Fe(II) | 1799 μM |
| Mn(II) | 911 μM |
| Cu(II) | 157 μM |
| Co(II) | 63 μM |
| Ca(II) | 677 μM |
| Ni(II) | 118 μM |

